# Design and implementation of suspended drop crystallization

**DOI:** 10.1101/2023.03.28.534639

**Authors:** Cody Gillman, William J. Nicolas, Michael W. Martynowycz, Tamir Gonen

## Abstract

We have developed a novel crystal growth method known as suspended drop crystallization. Unlike traditional methods, this technique involves mixing protein and precipitant directly on an electron microscopy grid without any additional support layers. The grid is then suspended within a crystallization chamber which we designed, allowing for vapor diffusion to occur from both sides of the drop. A UV transparent window above and below the grid enables the monitoring of crystal growth via light, UV, or fluorescence microscopy. Once crystals have formed, the grid can be removed and utilized for x-ray crystallography or microcrystal electron diffraction (MicroED) directly without having to manipulate the crystals. To demonstrate the efficacy of this method, we grew crystals of the enzyme proteinase K and determined its structure by MicroED following FIB/SEM milling to render the sample thin enough for cryoEM. Suspended drop crystallization overcomes many of the challenges associated with sample preparation, providing an alternative workflow for crystals embedded in viscous media, sensitive to mechanical stress, and/or suffering from preferred orientation on EM grids.

## Introduction

Crystallography is a widely used technique for determining the structures of both small and large molecules such as proteins (McPherson & Gavira, 2014). Crystals, which possess repetitive structural patterns, are utilized in this approach (McPherson, 1985). When a coherent beam of x-rays or electrons is directed at a crystal, it is scattered in predictable ways that provide information about the underlying structure of the molecule in the crystal (Bragg, 1912). Over the past century, a number of crystal growth methods have been developed and refined, including liquid-liquid diffusion (Salemme, 1972), vapor diffusion using hanging or sitting drops(McPherson, 1989), and lipidic cubic phase (LCP) (Landau & Rosenbusch, 1996). Additionally, two-dimensional crystallization utilizing dialysis and growth through evaporation and concentration has also been explored and documented (Gonen *et al*., 2005; Henderson & Unwin, 1975; Schmidt-Krey, 2007).

Vapor diffusion is the most commonly employed method for protein crystallization (Chayen & Saridakis, 2008). In this method, the protein of interest is mixed with a mother liquor and placed either in a small well or on a glass support that hangs above the solution. The mixture is then sealed in a chamber with additional crystallization solution to allow for vapor diffusion. As the vapors form, the effective concentration of the protein increases, causing the drop to shrink. Under certain conditions, crystals may form, which are then detected using light, UV, or fluorescence. Several automated instruments have been developed for crystal detection. Hanging drops are typically used for soluble proteins in aqueous solution, while sitting drops are preferred for membrane proteins that may be in a solution with detergent and lipids. Various conditions are tested to optimize crystal growth, including pH, temperature, precipitants, and additives.

MicroED is a cryogenic electron microscopy (CryoEM) technique that utilizes electron diffraction to determine the three-dimensional structure of proteins, peptides, and small molecules in cryogenic conditions (Shi *et al*., 2013; Jones *et al*., 2018; Sawaya *et al*., 2016; Xu *et al*., 2019; Gruene *et al*., 2018). This method is suitable for crystals that are extremely small and typically invisible to the naked eye, with a size a billionth that required for x-ray crystallography (Mu *et al*., 2021; Nannenga & Gonen, 2019*a*). Once a crystal is obtained, it is transferred onto an electron microscopy grid using a pipette and rapidly frozen in liquid ethane. The sample is then placed in an electron microscope operating at liquid nitrogen temperatures to minimize radiation damage. The electron beam is focused in diffraction mode onto the crystal when it is identified, and MicroED data is collected on a fast camera while the stage is continuously rotating. X-ray data reduction software is utilized to process the MicroED data, and established procedures are employed to determine the structures (Hattne, Reyes, Nannenga, Shi, De La Cruz *et al*., 2015).

In certain cases, it is advisable to avoid transferring crystals onto an electron microscopy grid. Some protein crystals may be too delicate and have a large solvent fraction, which can result in damage during the transfer process and render them unsuitable for MicroED. Additionally, membrane protein crystals embedded in lipids, such as those formed through lipidic cubic phase crystallization, are highly susceptible to damage from physical manipulation. For these sensitive samples, new sample preparation techniques must be developed and optimized to ensure their suitability for MicroED.

Recent studies have demonstrated successful determination of structures for membrane proteins embedded in lipids using a novel approach for sample preparation (Martynowycz *et al*., 2021, 2020, 2023). The method utilizes a scanning electron microscope coupled with a focused ion beam (FIB/SEM) for sample preparation. In one example, the human adenosine receptor (A2_A_AR) was crystallized in LCP, and the crystal drop was transferred to an electron microscopy grid by blotting and rapid freezing in liquid ethane. The sample was too thick for visualization by a transmission electron microscope, so fluorescence was used to locate the nanocrystals within the lipid matrix. Correlative light-EM was then utilized to expose the crystals with the FIB for MicroED analyses, resulting in a high-resolution (2.0 Å) structure of the human receptor (Martynowycz *et al*., 2023). A similar approach was also used to determine the structure of a functional mutant of the mammalian voltage-dependent anion channel VDAC (Martynowycz *et al*., 2020).

Although the aforementioned sample preparation methods have been successful, they rely on the assumption that crystals are not damaged during the physical manipulation and transfer onto an electron microscopy grid. Additionally, certain crystals, especially those that resemble sheets, may exhibit a preferred orientation on the grid carbon support, which can limit the reciprocal space available for sampling. Given these challenges, there is a need to develop alternative approaches for sample preparation for MicroED, as well as for other imaging applications such as x-ray crystallography.

Here we used conceptual design and 3D printing to create a suspended drop crystallization setup. This is a novel approach for sample preparation for MicroED that eliminates the need for crystal transfer and physical manipulation, offering an alternative to traditional crystallization methods. The method involves allowing crystallization to occur directly on an EM grid without support, enabling both sides of the drop to be exposed for uniform vapor diffusion. The absence of support film on the grid eliminates preferred crystal orientations and enables complete reciprocal lattice sampling. Crystal growth can be monitored visually, and the entire crystallization drop can be plunge-frozen directly on the EM grid. The method was successfully demonstrated on proteinase K crystals, resulting in a 2.1 Å resolution structure. This approach may have potential for other imaging applications beyond MicroED.

## Results

### The 3D printed suspended drop screening tool

The suspended drop crystallization screening tool is a screw cap that can mount pre-clipped EM grids and suspend them over a well reservoir. The screw and mounting arms are made of a flexible rubber material made of thermoplastic polyurethane (TPU) that applies gentle pressure on the clipped EM grid without the risk of bending (Figure 1A). The screw also incorporates a clear glass coverslip that is securely tightened by a 3D printed plastic screw to create a viewing window. After dispensing sample onto a support-free EM grid, the suspended drop is sealed into an incubation chamber containing mother liquor (Figure 1B). Suspended crystallization drops can be monitored through the viewing window using light and fluorescent microscopy. A screening tray has also been designed and 3D printed, which can accommodate multiple incubation chambers for larger screening experiments (Figure 1C). When suspended drop crystals are identified, the screening tool is unscrewed from the well, tweezers are used to retrieve the grid, and the grid is rapidly plunged into liquid nitrogen or ethane without blotting (Figure 1D). For MicroED, FIB/SEM milling is performed prior to TEM imaging (Figure 1E). The suspended drop crystallization method can also be used directly for x-ray analysis by mounting the grid directly onto the goniometer.

**Figure 1.**
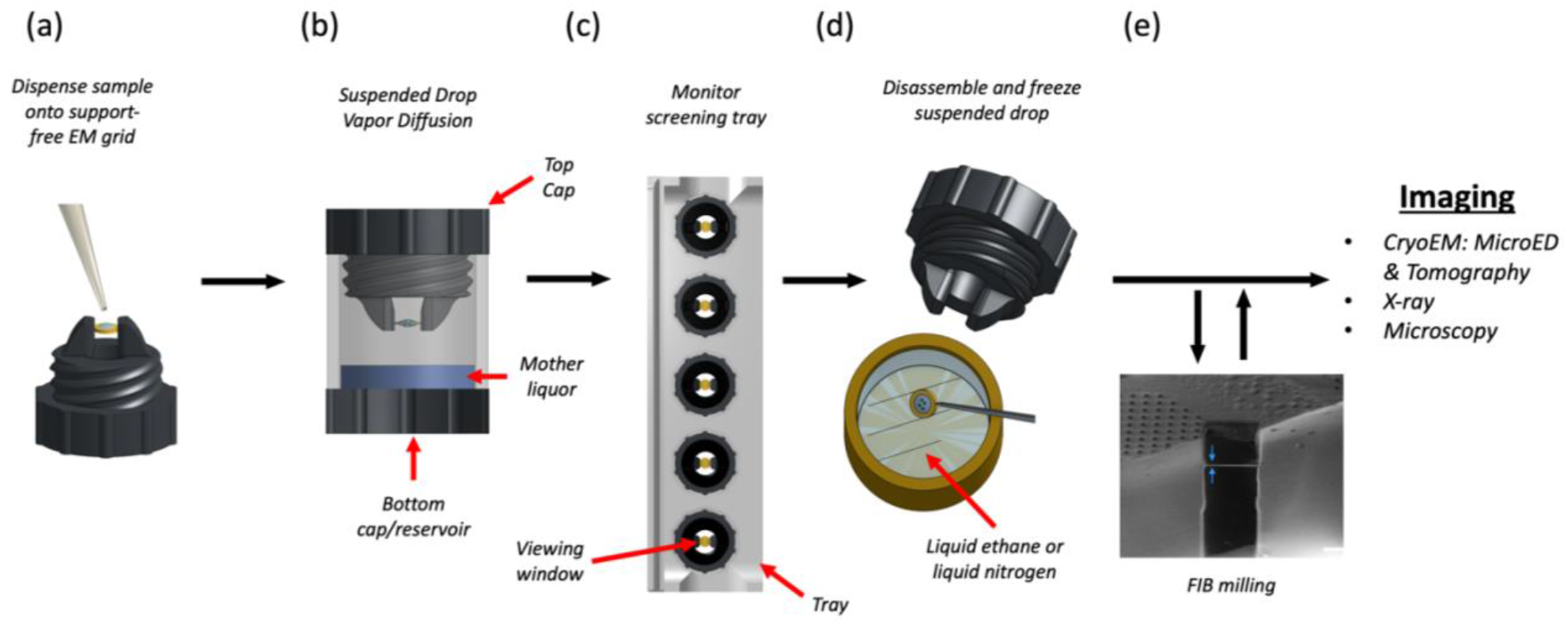
Suspended drop crystallization. (a) A support-free EM grid is clipped into an autogrid cartridge and mounted between the arms of the suspended drop screening tool. The sample and crystallization solution are dispensed onto the grid. (b) The chamber is immediately sealed to allow vapor diffusion. (b) The incubation chambers are inserted into a screening tray for efficient storing and monitoring of crystallization progress by light, fluorescence and UV microscopy. (d) EM grids containing crystals are retrieved from the screening tool and frozen. (e) The specimen is then interrogated by MicroED or other methods such as tomography, x-ray crystallography, or general microscopy. FIB milling is optional depending on the application.

### Protein crystals grown by suspended drop crystallization

We hypothesized that support-free gold gilder grids with a low mesh count (50-200 mesh) could be used to suspend crystallization drops during long incubation periods. Experimental results confirmed that suspended crystallization drops could be stably retained by such grids. To prepare the grids, 3 mm diameter gold gilder grids were clipped into autogrid cartridges for stability and rigidity, and glow-discharged before being mounted horizontally between the mounting arms of the screw cap. Proteinase K sample was mixed with mother liquor directly on the EM grid and the screening tool was tightened into the well of a crystallization tray for incubation (Figure 1A). Light microscopy and UV fluorescence was used to monitor the crystal growth through the coverslip at the top of the screening tool (Figure 2B, C).

**Figure 2.**
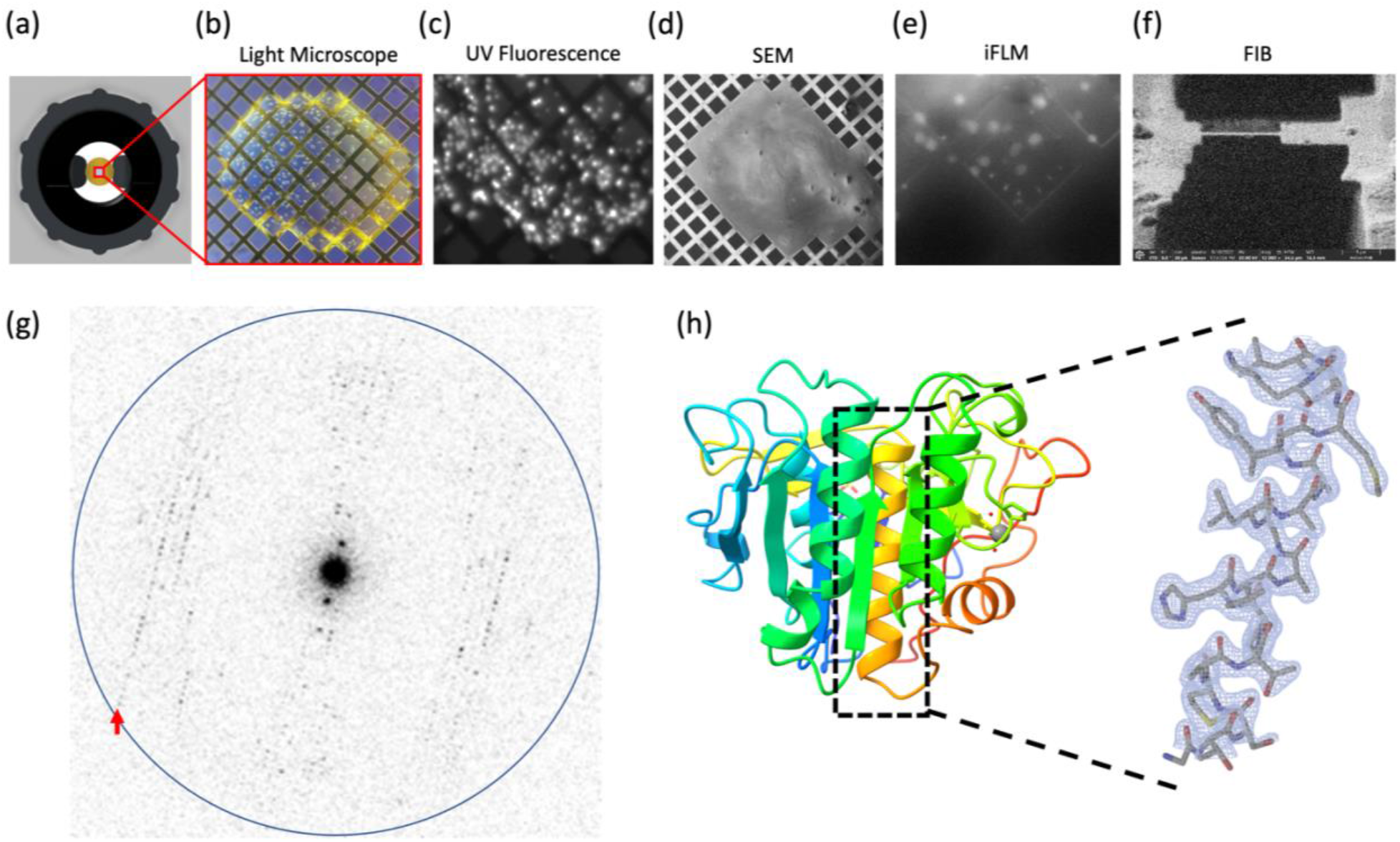
MicroED structure of suspended drop Proteinase K. (a) The suspended drop viewed from the top and imaged by (b) Light microscopy or (c) UV. A frozen suspended drop specimen was loaded into the FIB/SEM and imaged normal to the grid surface by (d) SEM and (e) iFLM with the 385 nm LED to locate submerged crystals. (f) The targeted crystal site was milled into a 300 nm thick lamella. (g) Example of MicroED data acquired from the crystal lamella. The highest resolution reflections visible to 2.1 Å (red arrow). Resolution ring is shown at 2.0 Å (blue). (H) Cartoon representation of the Proteinase K colored by rainbow with blue N terminus and red C terminus. The 2mFo–DFc map of a selected alpha-helix is highlighted, which was contoured at 1.5 σ with a 2-Å carve.

### Machining crystal lamella

Grids containing suspended proteinase K crystallization drops were retrieved from the screening apparatus and immediately plunged into liquid ethane. The grids were loaded into a plasma beam FIB/SEM equipped with an integrated fluorescence microscope (iFLM) at cryogenic conditions. The surface of the crystallization drop appeared smooth in the SEM and crystal features could not be observed (Figure 2D). To visualize crystals below the surface of the drop, the iFLM was used to detect crystal fluorescence (Figure 2E). A series of images was acquired at different focal points between the grid bars and the surface of the drop, and the depth at which the crystal appeared most in focus was taken to be the true depth of the crystal. The stack of reflective images was correlated to the X-Y plane of the SEM images and a three-dimensional representation of crystal locations inside the drop was generated.

The targeted crystal and surrounding media were milled into a thin lamella using a xenon plasma beam (Figure 2F). We used the xenon plasma beam because it is the fastest and most gentle option for milling crystals that are deeply embedded in solvent (Martynowycz *et al*., 2023). The final lamella was ∼7 μm wide and 300 nm thick.

### MicroED analyses of suspended drop crystals

The grid containing the crystal lamella was transferred to a cryogenically cooled Titan Krios electron microscope operating at 300 kV. The lamella site was identified with low magnification imaging and brought to eucentric height. A diffraction preview of the lamella was acquired to confirm that it would diffract to high-resolution (Figure 2G). Continuous rotation MicroED data was collected on a real space wedge from -40° to +40° tilt using a Falcon4 direct electron detector set to counting mode. Data was collected with a selected area aperture to reduce background dose. Strong and sharp reflections were visible to 2.1 Å resolution and a clear lattice was visible.

MicroED data were converted to standard crystallographic formats using our online tools which are freely available (https://cryoem.ucla.edu/microed). The data were indexed and integrated in XDS to 2.1 Å resolution. Phases for the MicroED reflections were determined by molecular replacement. The space group was determined to be P 4_3_2_1_2 with a unit cell of (a, b, c) (Å) = (68.26, 68.26, 101.95) and (α, β, γ) (°) = (90, 90, 90). The structure was refined using electron scattering factors (Table 1). The structure of proteinase K that was determined matches other MicroED structures of this protein that determined from crystals that were handled using traditional MicroED sample preparation protocols (Figure 2H).

**Table 1.**
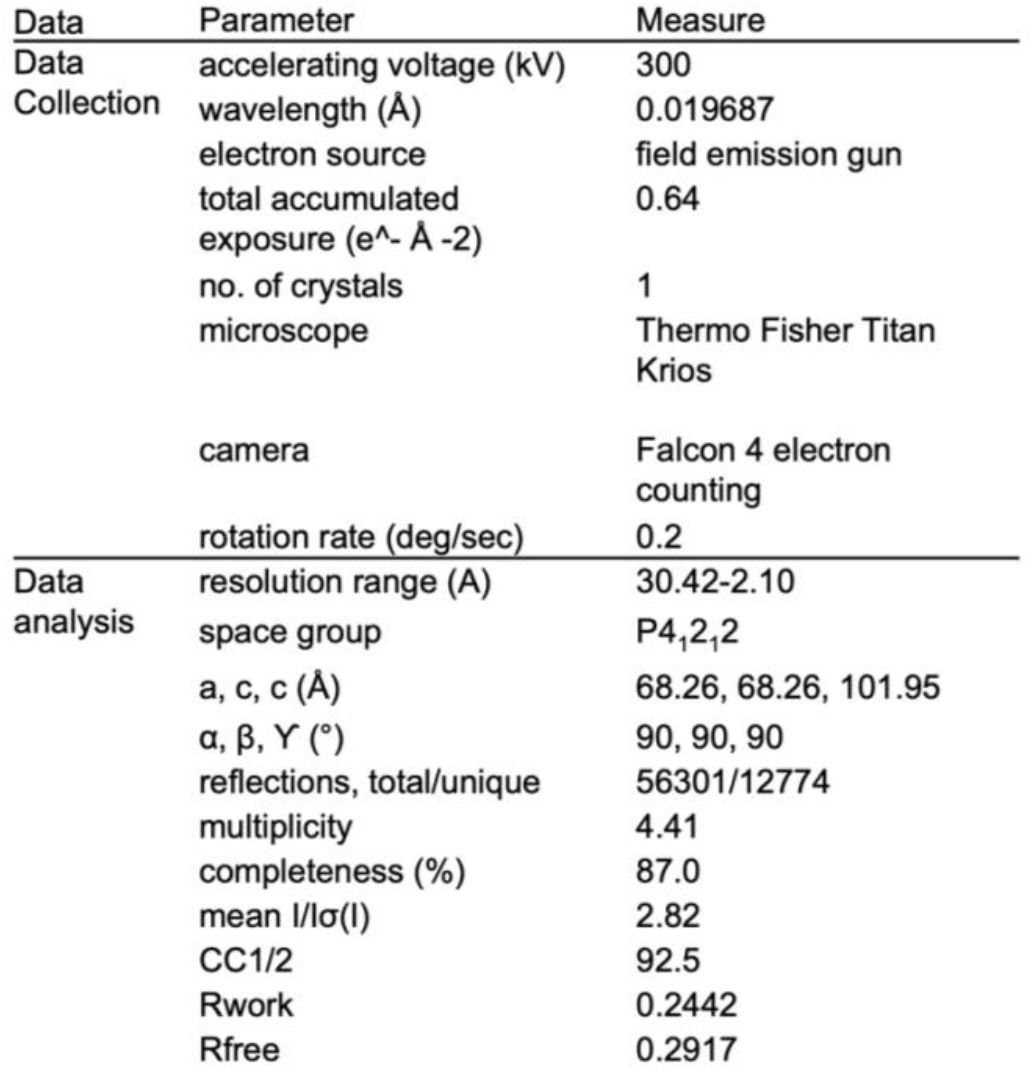
MicroED structure statistics of proteinase K crystallized by suspended drop.

## Discussion

In this study, we utilized suspended drop crystallization to grow crystals of a protein, and subsequently determined its structure by MicroED. To optimize the conditions for suspended drop crystallization, we developed a screening tool that features a screw cap with two extended arms for clamping an EM grid and a clear glass coverslip that creates a viewing window (see Figure 1A). Once the sample was dispensed onto a support-free EM grid, the suspended drop was sealed into an incubation chamber with mother liquor (see Figure 1B), and its growth could be monitored using light and fluorescent microscopy. To harvest the crystals, the screening tool was unscrewed from the incubation well, and the grid was retrieved with tweezers and rapidly frozen in liquid nitrogen or ethane for cryo-preservation.

The process for preparing MicroED samples is akin to the standard procedure followed in other cryoEM techniques like single particle analysis (SPA) and tomography. The sample is usually dispensed onto an EM grid, excess solvent is blotted, and the grid is vitrified by immersing it in liquid ethane (Nannenga & Gonen, 2019*b*). However, enhancing the preparation of samples for MicroED experiments using this method can be challenging because of limited options for improving crystal transfer and blotting conditions, which could cause damage to fragile crystals. Nevertheless, suspended drop crystallization offers an alternative specimen preparation method that eliminates the need for crystal transfer and blotting. This technique presents a promising solution for crystallographers dealing with challenging crystals that are embedded in viscous buffer (e.g., membrane proteins or crystals in high precipitant conditions), prone to mechanical stress, toxic, volatile, or limited in number in the drop. We envisage that suspended drop crystallization will be valuable in the preparation of recalcitrant crystals for MicroED experiments.

Crystals that adopt a preferred orientation on EM grids with carbon support can lead to incomplete sampling of the reciprocal space, limiting the accuracy of structural determination. This is especially common in plate-like sheet crystals, such as those of Catalase and Calcium-ATPase (Nannenga *et al*., 2014; Yonekura *et al*., 2015). However, growing crystals using the support-free suspended grid method can avoid preferred orientation. As there is no support film, crystals cannot align themselves in a specific orientation, allowing for 100% sampling of the reciprocal space for any crystal morphology and symmetry by merging data from several crystals. This approach is particularly useful for crystallographers working with challenging samples, enabling high-quality data collection and accurate structural determination.

The suspended drop crystallization tools described in this study enable crystal growth to be assayed in a sparse matrix directly on grids without support. The modular design allows for a large number of crystallization conditions to be assayed. While others have attempted to grow crystals directly on grids, they typically use carbon support and cannot perform sparse matrix crystallization assays (Li *et al*., 2018). We found that gold grids were the most inert and produced the most consistent results, as copper grids tend to oxidize and prevent crystal growth, and holey carbon grids can make it difficult to monitor crystal growth. Using support-free gold grids with a lower mesh count allows for easier monitoring of crystal growth, reduces the amount of material in contact with the sample, and decreases the likelihood of obstruction by grid bars, which is important for subsequent FIB milling. Additionally, because no blotting is required with this setup, the initial position of the crystals remains unchanged after freezing, which facilitates targeting and FIB milling.

Using a support-free grid and a sparse matrix approach, the suspended drop method allows for easier monitoring of crystal growth and eliminates physical contact with the sample. Additionally, the use of cryogenic plasma beam FIB/SEM enables efficient generation of sample lamellae, while cryogenic TEM allows for high-quality MicroED data collection. One major advantage of this approach is its potential applicability to membrane proteins, which are notoriously difficult to crystallize due to their softness and fragility. Furthermore, the use of automation and robotics could further streamline the process and make it more accessible to structural biologists. Overall, the suspended drop crystallization method has the potential to become a routine approach in structural biology and although not demonstrated in this study, suspended drop crystallization could be employed in x-ray crystallography, as well as other microscopy and cryoEM applications.

## Acknowledgments

This study was supported by the National Institutes of Health P41GM136508 and the Department of Defense HDTRA1-21-1-0004. The Gonen laboratory is supported by funds from the Howard Hughes Medical Institute. Coordinates and maps were deposited in the protein data bank (Accession code XXXX) and the EM Data bank (Accession code YYYY).

## Methods and Materials

### Materials

Proteinase K from *Tritirachium album* was purchased from Fisher BioReagents (Hillsborough, OR) and used without further purification. Ammonium sulfate and Tris buffer were purchased from Sigma-Aldrich (St. Louis, MO). All reagents were made with MilliQ water. The Ultimaker S5 3D printer and all filaments were purchased from MatterHackers (Lake Forest, CA). Glass coverslips were purchased from Ted Pella (Redding, CA). The gold gilder grids were purchased from Electron Microscopy Sciences (Hatfield, PA). The 100 nm fluorescent TetraSpeck Microspheres were purchased from Thermo-Fisher.

### Object design and 3D printing

All components of the screening tool were designed in the cloud-based CAD program Onshape.com and exported in STL file format. To generate GCODE files for 3D printing, the STL files were imported into the slicer program Ultimaker Cura 5.0 and default Ultimaker material profiles were used. All objects were printed at 0.1 mm layer height and 40 mm/sec print speed. All 3D printing was performed on an Ultimaker S5 3D printer equipped with a 0.4 mm diameter nozzle and a glass build surface with a layer of glue applied. The main body of the screening tool and the cover slip gasket were printed in thermoplastic polyurethane (TPU). The cover slip retaining screw was printed in co-polyester (CPE). A single on-grid screening tool takes approximately 1.5 hrs to print.

### Suspended drop crystallization

Proteinase K was dissolved in 0.1 M Tris-HCl pH 8.0 at 25 mg/ml. A support-free gold gilder grid was clipped into an autogrid cartridge, negatively glow-discharged for 1 min at 15 mA, and mounted in the screening tool. Equal volumes of proteinase K and 1.5 M ammonium sulfate were mixed dispensed on the mounted grid (∼0.3 µL final drop volume). The screening tool (with mounted grid and crystallization drop) was immediately screwed into a well of the crystallization tray containing 300 μL of 1.5 M ammonium sulfate in the reservoir. Within 48 hrs, crystals of Proteinase K were observed in the hanging crystal drops.

### Sample preparation and cryo-preservation

The EM grids supporting crystal drops were carefully removed from the screening tool with tweezers and rapidly plunged into liquid ethane. The grids were stored in liquid nitrogen until use.

### Machining proteinase K crystal lamellae using the plasma beam FIB/SEM

The vitrified EM grid was loaded into a Thermo-Fisher Helios Hydra dual-beam plasma beam FIB/SEM operating at cryogenic temperature. A whole-grid atlas of the drop was acquired by the SEM operating at an accelerating voltage of 0.5 kV and beam current of 13 pA using the MAPS v3.19 software (Thermo-Fisher). The crystal drop was coated with platinum by beam-assisted (argon beam at 4 nA, 5 kV) GIS coating for 1 min to protect the sample from ion and electron beams. The drop was then inspected using the iFLM with the 385 nm LED to locate crystals inside the drop at various Z dimensions. The sample was presented normal to the FIB beam and small holes were milled straight down the sample, around the crystal of interest, with the Xenon beam at 4nA for use as “fiducials” for the later correlation step. A comprehensive fluorescence stack of the crystals of interest was acquired with a binning of 2 (pixel size of 240 nm) and a step of 0.5 um (Figure 3B, top panel). This stack was deconvolved using the DeconvolveLab Fiji plugin (Sage *et al*., 2017). An experimental Point Spread Function (PSF) was measured using sub-resolution 100nm TetraSpeck microspheres. The processed PSF used for deconvolution was generated with the Huygens software (https://svi.nl/Huygens-Software). Further preprocessing using the 3D-Correlation Tool (3DCT) (Heymann *et al*., 2006) was performed: 1-stack reslicing in order to output isometric voxels 240 × 240 × 240 nm, and 2-intensity normalization. Low current FIB (10pA) and low voltage SEM (2 kV) images were acquired at grazing incidence (milling angle 11°) and were used to correlate against the fluorescent stack. 3DCT was used to correlate the SEM/FIB views with the fluorescence images. To do so, the milled holes were located in 2D in the SEM/FIB image and in 3D in the fluorescent stack. In the latter, the crystals of interest were located by delineating them with markers. In our hands, as low as 6 fiducial holes, both visible in fluorescence and SEM/FIB, were enough to correlate the two modalities with an error no less than 5 pixels.

This correlation process was performed during the milling procedure to make sure the final lamellae were on target. During the final steps of milling (when the lamella was a 2-3 μm thick), the correlation precision in Z was no longer enough. Milling was performed from top to bottom and the stage was brought back normal to the E-beam for checking the presence of the crystal at the surface of the lamella. To do so, SEM settings were set to 1.2 kV, 13 pA. These settings allowed scattering contrast between the crystal and the surrounding aqueous solvent. When the contours of the crystal were visible, milling was performed from bottom to top until the final thickness of 300 nm was reached. The xenon plasma beam (30 kV) was used for lamella milling at an angle of 11°. For the first milling step, two boxes (20 × 35 μm) separated by 5 μm 4 nA,. Second milling step a current of 1 nA was used to thin down the lamella to 3.5 μm. Third milling step a current of 0.3 nA was used to narrow the lamella to 10 μm wide (X dimension of the milling boxes) and thin it down to 2 um. Fourth milling step a current of 0.1 nA was used to thin down to 1 μm. Final milling step a current of 30 pA was used to generate a 300 nm thick lamella. The final lamella was 10 μm wide, 20 μm long, and 200 nm thick.

### MicroED Data Collection

Grids with milled lamellae were transferred to a cryogenically cooled Thermo-Fisher Scientific Titan Krios G3i TEM. The Krios was equipped with a field emission gun and a Falcon4 direct electron detector, and was operated at an accelerating voltage of 300 kV. A low magnification atlas of the grid was acquired using EPU (Thermo-Fisher) to locate milled lamellae. The stage was translated to the lamellae position and the eucentric height was set. The 100 µm selected area aperture was inserted and centered on the crystal to block background reflections. In diffraction mode, the beam was defined using a 50 μm C2 aperture, a spotsize of 11, and a beam diameter of 20 µm. MicroED data were collected by continuously rotating the stage at 0.2 ° / s for 400 s, resulting in a rotation range of 80°.

### MicroED data processing

Movies in MRC format were converted to SMV format using MicroED tools (Martynowycz *et al*., 2019; Hattne, Reyes, Nannenga, Shi, Cruz *et al*., 2015). The diffraction dataset was indexed and integrated in *XDS* (Kabsch, 2010*b*). Integrated intensities from a single crystal were scaled and merged in *XSCALE* (Kabsch, 2010*a*).

### Structure solution and refinement

Phases for the MicroED reflections were determined by molecular replacement in PHASER using Protein Data Bank (PDB) 6CL7 as the search model (McCoy *et al*., 2007; Hattne *et al*., 2018). The solution was space group P4_3_2_1_2 and unit cell dimensions 68.26, 68.26, 101.95 (a, b, c) (Å) and 90, 90, 90 (α, β, γ) (°). The first refinement was performed with Coot and phenix.refine (Afonine *et al*., 2012) using isotropic B-factors, automatic water picking, and electron scattering factors. Occupancies were refined for alternative side chain conformations and SO_4_ and calcium were placed in coordination sites. The final refinement used anisotropic B-factors, automatic water picking, and electron scattering factors and resulted in Rwork/Rfree = 0.2442/0.2917 and resolution of 2.1 Å.

## Notes

### Competing Interest Statement

The authors have declared no competing interest.

